# GenBio-PathFM: A State-of-the-Art Foundation Model for Histopathology

**DOI:** 10.64898/2026.03.17.712534

**Authors:** Saarthak Kapse, Mehmet Aygün, Elijah Cole, Emma Lundberg, Le Song, Eric P. Xing

## Abstract

Recent advancements in histopathology foundation models (FMs) have largely been driven by scaling the training data, often utilizing massive proprietary datasets. However, the long-tailed distribution of morphological features in whole-slide images (WSIs) makes simple scaling inefficient, as common morphologies dominate the learning signal. We introduce GenBio-PathFM, a 1.1B-parameter FM that achieves state-of-the-art performance on public benchmarks while using a fraction of the training data required by current leading models. The efficiency of GenBio-PathFM is underpinned by two primary innovations: an automated data curation pipeline that prioritizes morphological diversity and a novel dual-stage learning strategy which we term JEDI (**JE**PA + **DI**NO). Across the THUNDER, HEST, and PathoROB benchmarks, GenBio-PathFM demonstrates state-of-the-art accuracy and robustness. GenBio-PathFM is the strongest open-weight model to date and the only state-of-the-art model trained exclusively on public data.

## 1 Introduction

Modern histopathology requires the integration of complex, multi-scale morphological signals to perform tasks ranging from tumor subtyping and grading to molecular state inference [1–4]. While computational foundation models have emerged as a powerful solution for these tasks, their development has traditionally relied on a “brute-force” scaling paradigm using hundreds of millions of tissue tiles extracted from massive, often proprietary, whole-slide image (WSI) collections [5]. This approach is fundamentally constrained by the long-tailed distribution of pathology data, where common morphologies dominate training signals, while diagnostically critical features (e.g. rare cellular variants or transition zones) remain underrepresented [6]. In addition, these models are overwhelmingly based on the DINO family of models [7–9], leaving a notable gap in the exploration of new pretraining ideas.

Here we introduce GenBio-PathFM, a 1.1-billion-parameter foundation model that achieves state-of-the-art results across clinical, molecular, and robustness benchmarks (THUNDER [1], HEST [2], PathoROB [10]) while utilizing only 10–20% of the training data required by current leading models. The performance of GenBio-PathFM is driven by two innovations illustrated in Figure 1:

1. **Automated Data Curation:** A pipeline that identifies informative tiles for training. This “quality over quantity” approach ensures the model focuses on morphological diversity rather than redundant patterns. Full details can be found in Section 4.1.
2. **JEDI (JEPA + DINO) Pretraining:** A novel dual-stage pretraining recipe that transitions from coarse discriminative pretraining to a fine-grained masked image modeling stage. Full details can be found in Section 4.4.

**Fig. 1:**
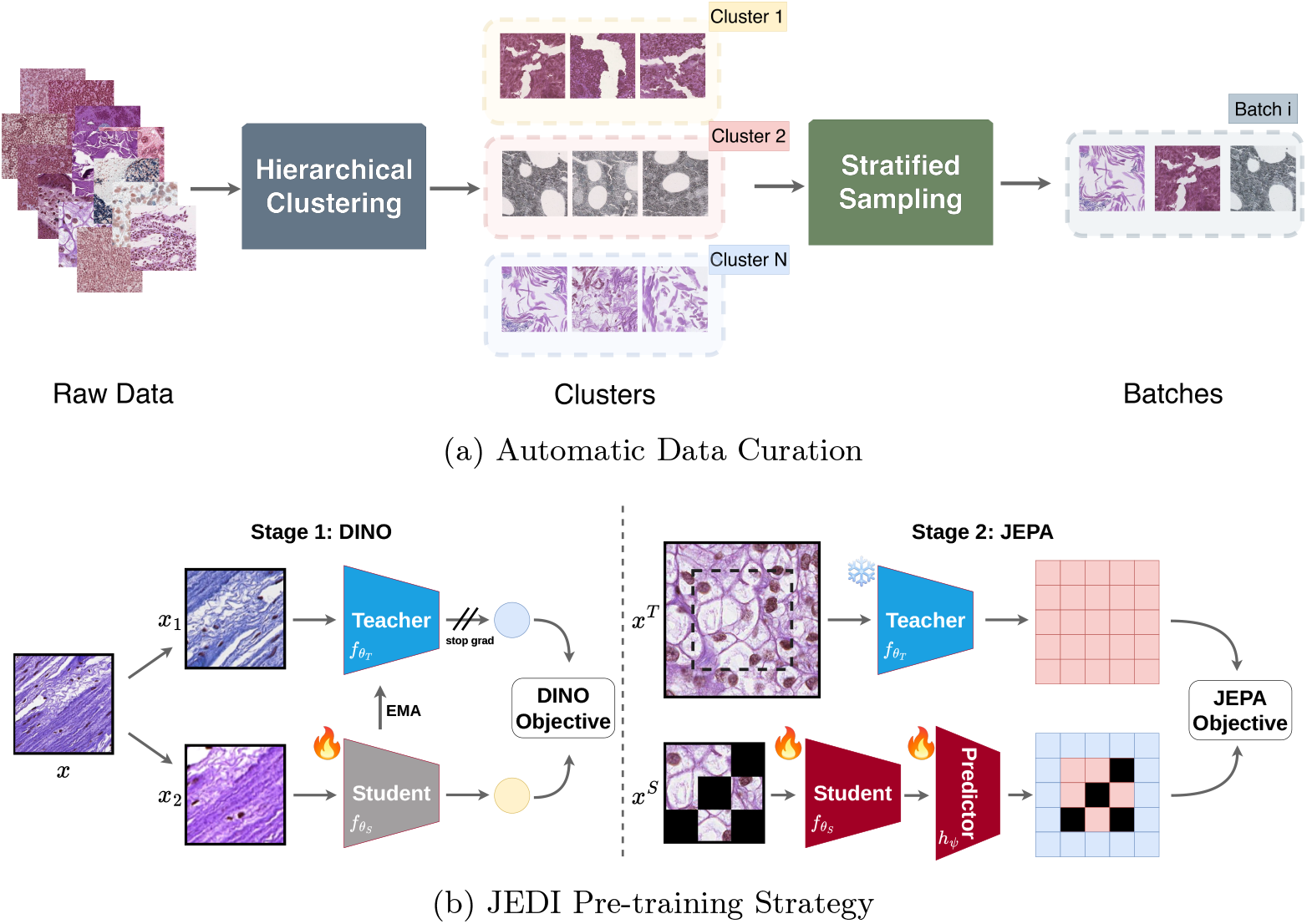
Overview of the GenBio-PathFM data processing and training framework. (a) Our automatic curation pipeline utilizes hierarchical clustering and stratified sampling to select morphologically diverse training data. (b) The JEDI pretraining strategy follows a two-stage approach: DINO-based self-supervised training (Stage 1) followed by a JEPA objective (Stage 2) which augments standard masked region prediction with visible region prediction and outpainting. For Stage 2, both the teacher and student networks are initialized with the weights of the Stage 1 teacher, after which the Stage 2 teacher is kept completely frozen. Note that we illustrate Stage 1 for one single pair of views (*x*_1_, *x*_2_) and omitted projection heads for simplicity.

In the first stage of JEDI, we train a DINO-based [7–9] model to capture robust, global morphological features. In the second stage, we freeze this encoder and use it as a teacher for a Joint-Embedding Predictive Architecture (JEPA)-based [11] student tasked with inpainting and outpainting masked regions. This second stage is intended to encourage the model to develop fine-grained spatially-aware representations. While recent video modeling work has shown that MAE-pretrained frozen teachers can guide JEPA-style predictive learning [12], GenBio-PathFM is, to our knowledge, the first to establish this paradigm for static images and the first to employ a teacher pretrained with a view-invariance objective.

By combining intelligent data curation with a novel pretraining recipe, GenBio-PathFM establishes that high-performance computational pathology can be achieved with dramatic improvements in data and compute efficiency. This work provides a scalable, robust, and fully open-weight foundation for clinical diagnostic assistance and biological discovery.

## 2 Results

### 2.1 GenBio-PathFM achieves state-of-the-art performance with unprecedented data efficiency

Recent histopathology foundation models have relied on increasingly large proprietary datasets to improve performance [3, 4, 13]. We demonstrate that GenBio-PathFM learns competitive representations using significantly less data. We evaluate frozen GenBio-PathFM on two public pathology benchmarks: THUNDER [1] and HEST [2]. The results of these comparisons are summarized in Figure 2. We see that GenBio-PathFM achieves performance similar to or better than prior open-weight models on both benchmarks. GenBio-PathFM outperforms all other models on the THUNDER benchmark. On the HEST benchmark, GenBio-PathFM performs on par with H-Optimus-1 (0.420 vs. 0.422) despite more than 5-fold reduction in training data (18% as many WSIs). On HEST, GenBio-PathFM is the top performer on more subtasks than any other model (Table A1); meanwhile, on THUNDER, it ties with H-Optimus-1, with both models ranking first in 3 out of 12 subtasks (Table A3). These findings show that it is possible to achieve state-of-the-art performance without the need for massive private data repositories.

**Fig. 2:**
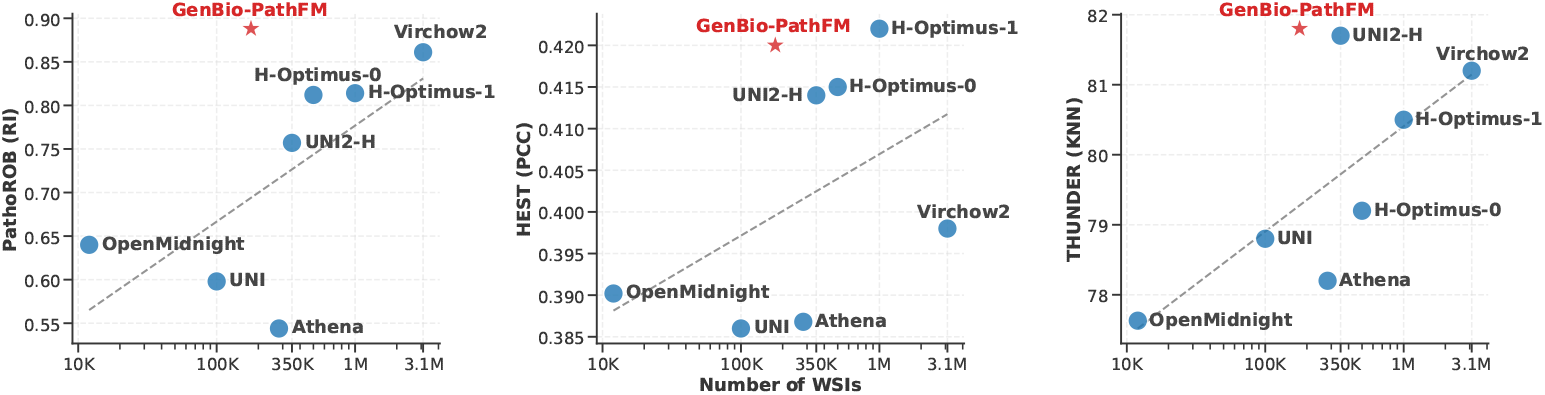
Data efficiency of histopathology foundation models across three benchmarks: PathoROB (left), HEST (center), and THUNDER (right). The dashed line represents the trend for all models before GenBio-PathFM.

We also investigated the model’s ability to transfer across diverse organ types and cancer contexts by training a single predictor jointly across all available HEST subtasks. This yielded a significant performance boost, increasing the average Pearson correlation from 0.420 to 0.669 (Table A2). This capability demonstrates that GenBio-PathFM serves as a highly expressive foundation for complex tasks like prediction of gene expression across multiple tissues.

### 2.2 GenBio-PathFM sets a new state-of-the-art for robustness to technical variation

To assess clinical reliability, we evaluated GenBio-PathFM’s resilience to non-biological technical variations (e.g. staining protocols, scanner hardware) using the PathoROB benchmark. Across three multi-center datasets (Camelyon, TCGA, Tolkach ESCA), GenBio-PathFM established a new state-of-the-art average Robustness Index (RI) of 0.888, surpassing established baselines including Virchow 2 (0.861) and UNI2-H (0.757) (Table A4). This robustness was particularly pronounced on the Camelyon dataset; despite high inter-center variability, GenBio-PathFM maintained an RI of 0.865, significantly outperforming UNI2-H (RI = 0.544) despite both models achieving high balanced accuracy (BACC).

We further assessed the performance decay between in-distribution (ID) and out-of-distribution (OOD) data to assess robustness to distribution shift. GenBio-PathFM exhibited exceptional stability, with a much lower average ID performance drop (1.0) compared to H-optimus-1 (1.9) and UNI2-H (2.6) (Table A5), and no average performance drop on OOD data. In the Tolkach ESCA cohort, only GenBio-PathFM achieved no performance drop on ID data, suggesting that its learned representations are largely invariant to site-specific noise.

When mapping the trade-off between downstream task performance and robustness (Figure A1), GenBio-PathFM occupies the upper-right frontier of the design space, achieving a high average BACC (96.9%) without the robustness drops of current state-of-the-art models.

### 2.3 GenBio-PathFM exhibits balanced generalization across clinical and molecular domains

Beyond individual benchmarks, we assessed the holistic performance of GenBio-PathFM across the THUNDER (clinical subtyping/grading), HEST (spatial transcriptomics), and PathoROB (technical robustness) frameworks. Unlike existing models that often exhibit “peak” performance on specific tasks at the expense of others, GenBio-PathFM demonstrates a consistently high performance profile across all three axes (Fig. 2).

Our analysis indicates that GenBio-PathFM maintains competitive accuracy across heterogeneous clinical and biological objectives without pronounced weaknesses. For instance, while some models prioritize morphological grading (THUNDER), they often show diminished capacity for morphology–expression mapping (HEST). In contrast, GenBio-PathFM achieves state-of-the-art results in clinical subtyping while simultaneously maintaining high correlation with gene expression patterns and resilience to cross-site technical variation (Fig. 3). This stable behavior across diverse histopathology tasks suggests that GenBio-PathFM has learned a more “universal” representation of tissue morphology, establishing it as a highly versatile foundation model for broad clinical and research applications.

**Fig. 3:**
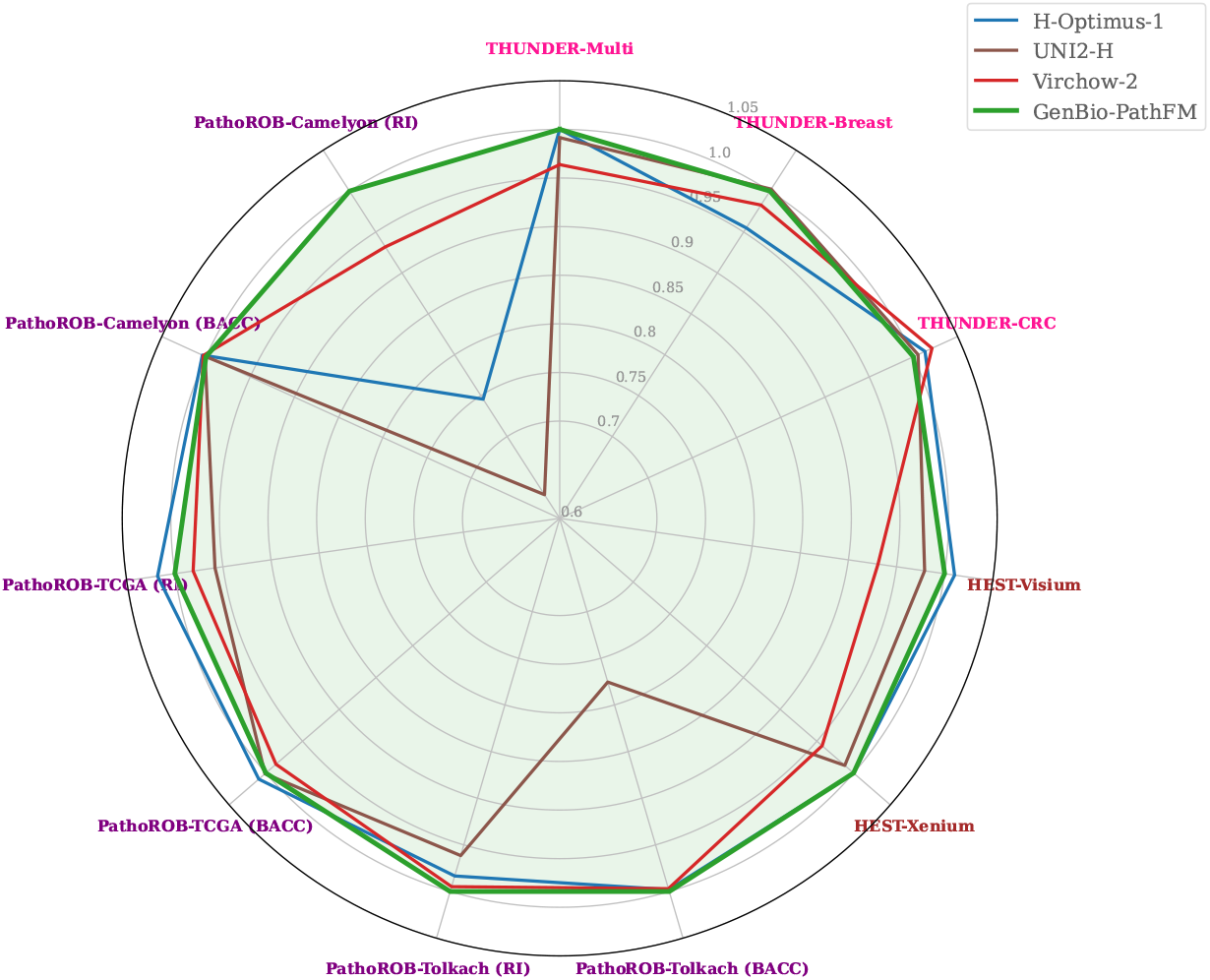
Performance of open-weight histopathology foundation models. The 11 tasks are drawn from the THUNDER, HEST, and PathoROB benchmarks. All scores are normalized to the performance of GenBio-PathFM, which is represented by the 1.0 radial line.

## 3 Discussion

Our results suggest that the performance gains in histopathology foundation models, previously attributed primarily to dataset scaling, can be achieved through targeted architectural and training refinements. By reaching performance parity with H-Optimus-1 on the HEST benchmark (using 18% of the training data) and out-performing Virchow2 on robustness (using only 6% of its training data) (Fig. 2), GenBio-PathFM demonstrates that high-quality representation learning is not strictly dependent on massive, proprietary data volume. These findings imply that the field may be entering a phase of diminishing returns for “brute-force” scaling, shifting the focus toward strategic data selection and representation learning to drive model utility.

Specifically, this study supports a paradigm where “quality-first” data curation is fundamentally coupled with advanced pretraining objectives. Because histopathology data is highly redundant, our curation pipeline prioritizes “high-entropy” content—such as rare histological variants and complex tissue architectures—over the unconstrained aggregation of slides. Notably, our pretraining data is highly skewed: over 54,000 of the approximately 177,000 WSIs correspond exclusively to skin tissue, predominantly from the HistAI cohort. Consequently, the available WSI count for all other organs is low. Despite this, GenBio-PathFM achieves state-of-the-art generalization on downstream tasks involving underrepresented organs. This robust performance further underscores the extreme data efficiency of our curation pipeline. Furthermore, while standard DINO pretraining establishes a strong morphological baseline, we advance this paradigm to extract richer, fine-grained spatial understanding from our diverse curated data. Our novel JEDI strategy accomplishes this by building upon the initial view-invariant feature learning (Stage 1) with a dedicated predictive feature-space objective (Stage 2). The explicit benefit of this second stage is demonstrated in our ablation studies (Table A6), where the addition of the JEPA objective yields substantial gains in both robustness (PathoROB increases from 0.838 to 0.888) and expression inference (HEST increases from 0.410 to 0.420) over Stage 1 alone. By combining morphological diversity with these optimized learning objectives, GenBio-PathFM captures a more biologically meaningful representation of tissue, evidenced by its stability in out-of-distribution scenarios on the PathoROB benchmark (Table A5).

The balanced performance across clinical, molecular, and robustness benchmarks (Fig. 3) further positions GenBio-PathFM as a versatile backbone for multimodal applications. Unlike models that exhibit trade-offs between specific diagnostic tasks, GenBio-PathFM exhibits strong performance across heterogeneous objectives. The significant accuracy boost observed during joint training across HEST tasks (Table A2) highlights the expressiveness of these representations.

There are several limitations to this work. First, we restrict our primary analysis to open-weight models. While some proprietary models [14, 15] report competitive benchmark scores, such models cannot be built upon by the community or independently validated. Second, while GenBio-PathFM is highly data-efficient, the computational resources required for its pretraining are still non-trivial. Finally, further research is needed to determine how these efficiency gains translate to even larger data regimes and more diverse multimodal inputs.

In summary, GenBio-PathFM demonstrates that intelligent data curation and optimized learning objectives can effectively complement or even exceed the gains of unconstrained data scaling. By providing an open-weight foundation model that is both robust and data efficient, we offer a framework for the development of pathology AI that is more accessible, transparent, and biologically useful.

## 4 Methods

### 4.1 Pretraining Data

#### 4.1.1 Data Sources

Our pretraining corpus comprises approximately 177k Whole Slide Images (WSIs) aggregated from four open-access sources (Table B8). HistAI [16] provides the plurality of our data and covers two individual organs and seven organ systems. This is supplemented by TCGA [17], ensuring broad coverage of 32 distinct malignancy types. To increase the coverage of non-pathological morphology, we incorporate WSIs from 23 normal tissue types via the GTEx portal [18]. Finally, WSIs from the REG challenge [19] are included to capture the morphological transitions between healthy, precancerous, and cancerous states across 7 organs. Table B9 compares the scale and accessibility of our data to that of current SOTA models.

#### 4.1.2 Preprocessing

This section described how we use our WSI collection to produce sets of tiles for training.

##### 1. Global region of interest identification

We begin with a collection of WSIs at a variety of magnifications (5×, 10×, 20×, 40×). For each WSI, we use the TRIDENT framework [20] (with default hyperparameters) to segment the area covered by tissue. This allows us to filter out any regions that are completely empty. Following [3], we then partition each WSI into tiles of size 392 × 392 and retain all tiles that have nonzero overlap with the tissue area.

##### 2. Tile-level filtering

Each pixel in each patch is classified as tissue or background by applying an HSV filter. To be precise, pixels are classified as tissue if

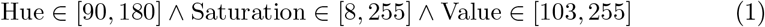

and otherwise they are classified as background. We discard all tiles consisting of more than 35% background pixels. The filtering criteria and their values are adopted from Virchow2 [3].

The final result is a set of tiles 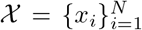, where *x*_*i*_ ∈ ℝ^392×392×3^ and *N* ≈ 8 × 10^8^. Note that 𝒳 is further processed using either unsupervised data curation (Section 4.2) or metadata-based data curation (Section 4.3) to produce training sets 𝒳_*S*1_ ⊂ 𝒳 (for pretraining stage 1) and 𝒳_*S*2_ ⊂ 𝒳 (for pretraining stage 2).

### 4.2 Unsupervised Data Curation

This section describes our fully unsupervised data curation pipeline, which selects a diverse subset of informative tiles without relying on manual annotations. Our approach is inspired by clustering-based data selection strategies for self-supervised learning [21, 22].

#### 1. Tile embedding

Let *v*_*ϕ*_ : ℝ^*H*×*W*×3^ → ℝ^*d*^ represent an image encoder parameterized by *ϕ*, where *d* is the image encoder’s embedding dimension. For each image *x*_*i*_ ∈ 𝒳, we compute an embedding *z*_*i*_ = *v*_*ϕ*_(*x*_*i*_). Let 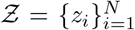. We implement *v*_*ϕ*_ as a pretrained CLIP ViT-B/32 encoder [23].

#### 2. Hierarchical clustering

We adopt the hierarchical clustering framework of [22]. We first perform a clustering step (using the embeddings Ƶ) that partitions 𝒳 into *K*_1_ basic concepts

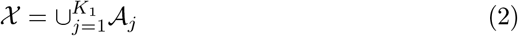

where 𝒜_*j*_ ∩𝒜_*k*_= ∅ for all *j* ≠ *k*. We then cluster these basic concepts into *K*_2_ < *K*_1_ higher-level concepts:

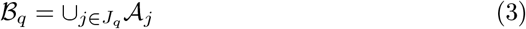

where 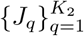 form a partition of {1, …, *K*_2_}. We set *K*_1_ = 10^5^ and *K*_2_ = 10^4^. We speculate that a deeper hierarchy would result in more clusters that correspond to artifacts (e.g. tissue folds, ink marks). We performed the full hierarchical clustering procedure separately for each magnification and combined the results. Hyperparameters are as follows: 50 centroid updates for each *K*-means step; 10 sampling steps for the resampling procedure at each hierarchy level.

#### 3. Stratified sampling

We construct 𝒳_S1_ by sampling from the concepts discovered during hierarchical clustering. We adopt a stochastic sampling scheme similar to that used in DINOv3 [9]. See Algorithm 1 for details. We set *p* = 0.9 and *t* = 4096. We define 𝒯_mag_ by the probabilities in Table B10.

We apply this pipeline to 𝒳 (Section 4.1.2) to produce 𝒳_*S*1_ ⊂ 𝒳, which is used in the first stage of pretraining. Our final 𝒳_*S*1_ consists of approximately 400M tiles.

##### Algorithm 1

Sampling procedure for 𝒳_S1_.

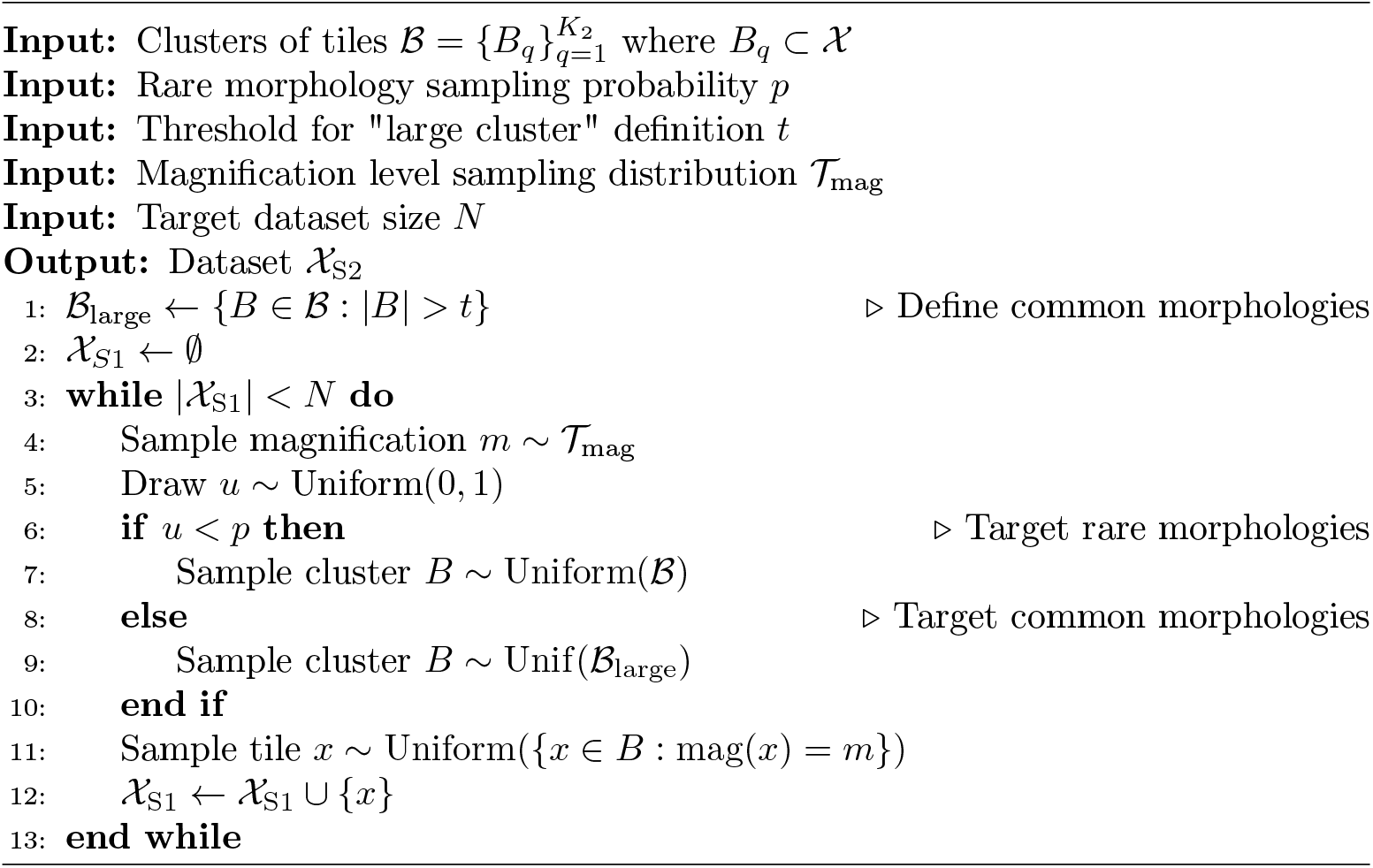

### 4.3 Metadata-based Data Curation

Here we describe the metadata-driven data curation pipeline [24], which selects tiles in an attempt to balance three WSI-level characteristics: magnification level, data source, and tissue type.

In what follows, we let *g* ∈ *G* represent a particular (magnification, data source, tissue type) combination, where *G* is the set of such combinations represented in 𝒳. Also let 𝒲_*g*_ ⊂ 𝒳 denote the collection of all available patches corresponding to a given metadata combination *g*.

#### 1. Intra-group clustering

To ensure morphological diversity within each metadata group, we cluster the tiles in 𝒲_*g*_ for each *g* ∈ *G*. Let *v*_*ϕ*_ : ℝ^*H*×*W*×3^ → ℝ^*d*^ represent an image encoder. For each patch *x*_*i*_ ∈ 𝒲_*g*_, we compute an embedding *z*_*i*_ = *v*_*ϕ*_(*x*_*i*_). We perform *K*-means clustering on the collection of embeddings {*z*_*i*_ } for this combination. Let 𝒞_*g*_ denote the partition of 𝒲_*g*_ produced by these clusters. We set *K* = 10 and used DINOv3-pretrained ViT-S/16 (21M) distilled model [9] as *v*_*ϕ*_.

#### 2. Sampling

We populate 𝒳_*S*2_ by following Algorithm 2. 𝒯_mag_ is defined in Table B10. We define 𝒯_src|*m*_ using Table B11. 𝒯_tissue|*m,s*_ is defined in Tables B12, B13, and B14.

We apply this pipeline to 𝒳 (Section 4.1.2) to produce 𝒳_*S*2_ ⊂ 𝒳, which is used in the second stage of pretraining. Our final *X*_*S*2_ consists of approximately 400M tiles.

##### Algorithm 2

Sampling procedure for 𝒳_*S*2_.

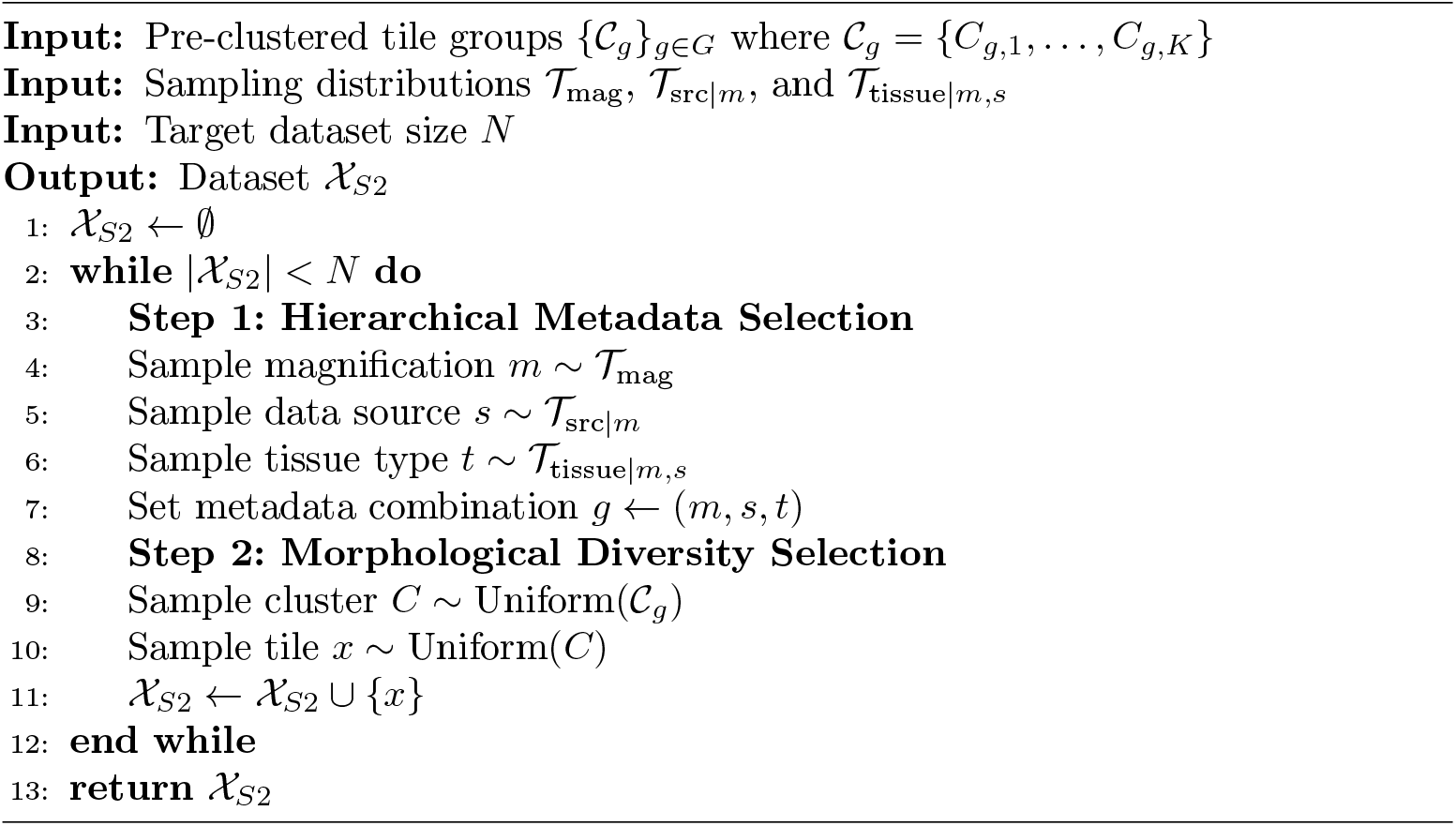

### 4.4 GenBio-PathFM

In Section 4.4.1 we describe the architecture of GenBio-PathFM. We introduce our two-stage pretraining strategy, which we call JEDI (**JE**PA + **DI**NO), in Section 4.4.2 and Section 4.4.3.

#### 4.4.1 Architecture

As before, let *x*_*i*_ ∈ ℝ^*H*×*W*×3^ be a single tile. Let *f*_*θ*_ : ℝ^*H*×*W*^→ ℝ^*q*^ be a *single-channel* feature extractor where *q* is the embedding dimension. To generate an embedding of *x*_*i*_, we first embed each channel

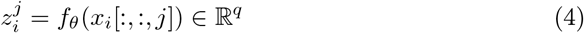

for *j* ∈ {0, 1, 2} and then form

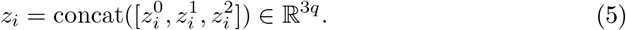

The tile embedding *z*_*i*_ can then be passed to prediction heads which feed into the loss function, whose details are described in Section 4.4.2 and 4.4.3.

This channel-agnostic late fusion formulation is intended to encourage the model to emphasize morphology over staining intensity, which can vary significantly between data collection sites.

We implement *f*_*θ*_ as a modified 1.1B-parameter ViT-G backbone based on DINOv3 [9]. The embedding dimension is *q* = 1536. We use 40 Transformer blocks with an embedding dimension of 1536 and 24 attention heads. We replace the standard learnable absolute positional embeddings with Rotary Position Embeddings (RoPE) [25]. Note that the ViT-G backbone in DINOv3 [9] uses four register tokens, which we include as well.

#### 4.4.2 JEDI Pretraining: Stage 1

We use a scalable self-supervised learning (SSL) framework based on the DINO family of models [8, 9]. The model utilizes a student-teacher setup to optimize two objectives: a global cross-entropy loss for feature consistency and a masked image modeling (MIM) loss for local spatial reasoning.

##### Architectural components

Write the teacher backbone as 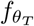 and the student backbone as 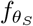. The backbone produces per-channel latent representations 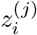 which are processed by two projection heads:

- **DINO head**. A 3-layer Multi-Layer Perceptron (MLP) with a hidden dimension of 2048 and a bottleneck dimension of 384, which maps the features to 131,072 prototypes. Denote these projection heads as 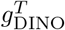 for the teacher and 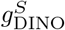 for the student.
- **MIM head**. A 3-layer MLP with a 2048 hidden dimension, a 256 bottleneck dimension, and an output space of 98,304 prototypes. Denote these projection heads as 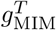 for the teacher and 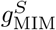 for the student.

Both heads produce temperature-scaled probability distributions. We apply a KoLeo regularizer [26], ℒ_*koleo*_, to the class tokens to encourage a uniform feature spread across the batch. We apply Sinkhorn-Knopp normalization to teacher projection head outputs.

##### Training objective

Let 𝒱_*global*_ and 𝒱_*local*_ represent the sets of unmasked global and local crops from an input tile. Let 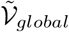 represent the global crops with masking applied. Define 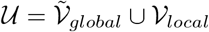.

The global DINO loss minimizes the cross-entropy *H* between projection outputs across all valid pairs, which are defined to be pairs (*x, x*^*′*^) where the student crop *x*^*′*^ does not originate from the same view as the teacher crop *x*, written as *x* ≠ *x*^*′*^. Then the loss is:

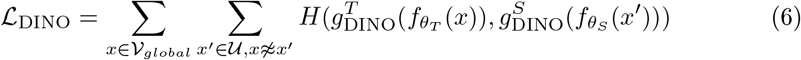

Now we turn to the MIM loss. For each global crop *x* ∈ 𝒱_*global*_, we define a masking operation mask(*x*, ℳ) where ℳ is a collection of indices indicating which patches to mask. This operation replaced patches at indices *i* ∈ ℳare replaced with a learnable mask token vector. The MIM objective minimizes the cross-entropy between the teacher’s patch predictions and the student’s reconstructed patch predictions, summed over the masked indices:

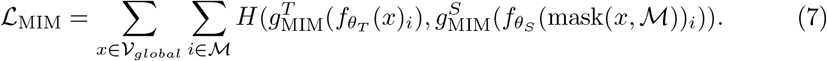

The total objective ℒ is the weighted sum of the global DINO cross-entropy loss, the dense MIM cross-entropy loss along with the koleo regularizer:

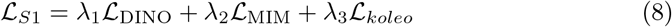

where we set *λ*_1_ = 1.0, *λ*_2_ = 0.5, *λ*_3_ = 0.1.

##### Augmentation details

We employ a multi-crop strategy, extracting global crops of size 224 × 224 and local crops of size 96 × 96 from the original 392 × 392 pixel patches. The data augmentation pipeline includes both geometric (random resized cropping and flipping) and semantic transformations from DINO. Specifically, to ensure the representations are robust to multi-site scanning and staining artifacts, we apply color jittering and Hematoxylin-Eosin-DAB (HED)-based stain perturbations [27].

##### Optimization details

We train our model using a Warmup-Stable-Decay (WSD) learning rate schedule [28]. Training is performed for a total of 150k iterations. This includes 10k iterations of warmup, followed by 100k iterations of stable learning, and a final 40k step decay phase to zero. The stable learning rate is set to 2 × 10^−4^. Optimization is performed using AdamW [29] with a weight decay of 0.04, *β*_1_ = 0.9, *β*_2_ = 0.99, and gradient clipping at 10.0 to prevent instability. We adopt a student-teacher momentum of 0.994. We use a batch size of 2048.

##### Multiscale post-training

To ensure robustness across resolutions, we include a post-training phase. We sample from the following pairs of global/local crop resolutions with the following probabilities: (224, 96) with *p* = 0.8, (256, 112) with *p* = 0.1, and (288, 128) with *p* = 0.1. This phase consists of 10k steps using a learning rate of 10^−5^ that decays to zero.

#### 4.4.3 JEDI Pretraining: Stage 2

In the second stage, we transition to a Joint-Embedding Predictive Architecture (JEPA) objective. The encoder from Stage 1 is frozen and used as a teacher 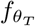. A student network 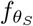 is initialized with the teacher’s weights and trained alongside a prediction head *h*_*ψ*_ to predict the teacher’s embeddings from partial inputs.

##### Predictor architecture

We implement the prediction head *h*_*ψ*_ as an asymmetric 8-block Vision Transformer decoder. The 1536-dimensional latent representations from 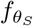 are first linearly projected to 512 dimensions. Learnable mask token ([MASK]) is repeated and then concatenated to represent missing regions, and Rotary Position Embeddings (RoPE) are applied. A final linear layer projects the result back to 1536 dimensions to align with the embedding space of 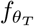.

##### Spatial masking scheme

Given a tile *x*, we define a teacher crop 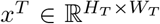 and a student crop 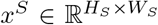. The student crop is a smaller center crop of the teacher crop, so *H*_*T*_ > *H*_*S*_ and *W*_*T*_ > *W*_*S*_. We partition the patch indices of *x*^*S*^ into a visible set 𝒱 and a masked set ℳ. The regions outside the student’s view *x*^*S*^ but within the teacher’s full extent *x*^*T*^ form the outpainting set 𝒪. We set *H*_*T*_ = *W*_*T*_ = 288 and *H*_*S*_ = *W*_*S*_ = 224.

##### Training procedure

The teacher encoder processes all patches to produce 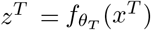. The student encoder processes only the visible patches to produce 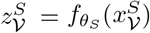. The prediction head *h*_*ψ*_ then takes these visible embeddings, concatenates them with the repeated learnable mask token ([MASK]) for the missing indices, and generates predictions 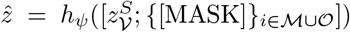 corresponding to all patches represented in *z*^*T*^, where *z*^*T*^ ∈ ℝ^*L*×*q*^ and 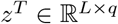, and *L* = | 𝒱 ∪ ℳ ∪ 𝒪 | denotes the total number of patches in the teacher crop. We then train to minimize

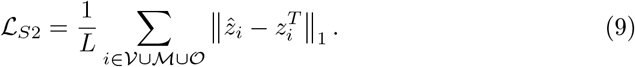

Note that register tokens and the CLS token are not used in the above calculation.

##### Augmentation details

We adopt a random 50% masking strategy for the student network input, similar to MAE, diverging from the multi-block-wise masking typical of JEPA. Data augmentations were limited to horizontal and vertical flipping, explicitly omitting color-space and HED perturbations. Furthermore, the random cropping scale was constrained to [0.7, 1.0] with aspect ratio [0.75, 1.33] to better align the training data with the visual distribution typically encountered during inference.

##### Optimization details

Training is performed for a total of 20k iterations, which includes 10k iterations of warmup. We use an effective batch size of 4096 distributed across 64 GPUs. The base learning rate is set to 1.5 × 10^−4^, resulting in an effective learning rate of 2.4 × 10^−3^ based on a linear scaling rule with respect to a base batch size of 256. Optimization is performed using AdamW with *β*_1_ = 0.9 and *β*_2_ = 0.95.

### 4.5 Evaluation

We evaluated our model on three public computational pathology benchmarks: THUNDER, HEST, and PathoROB. Unless otherwise noted, we used the CLS token as the representation and kept the network frozen, testing the generalization of the learned embeddings without additional fine-tuning. We followed the original code and protocols for each benchmark exactly to ensure a consistent and fair evaluation. THUNDER [1] provides a rigorous assessment across 12 diverse datasets spanning multiple organs, including breast, colorectal, and lung, as well as a variety of tissue types. We focused on the *k*-nearest neighbor (kNN) task to measure the model’s performance and its ability to encode broad histological structures. Performance on this benchmark is reported using F1-score.

HEST [2] evaluates a model’s capacity to link morphology to molecular phenotypes by predicting gene expression from histology. It comprises nine tasks across eight human cancers and nine organs, including primary sites such as breast (IDC), prostate (PRAD), and lung (LUAD), along with metastatic lymph nodes. Following the original benchmark protocol, we trained a single ridge regression model on principal components of the frozen embeddings to predict the top 50 high-variance genes. This setup tests sensitivity to subtle morphological cues associated with transcriptomic variation. Performance is measured via the Pearson correlation coefficient.

PathoROB [10] assesses model robustness using multi-center datasets and provides metrics such as the Robustness Index to determine whether learned representations capture true biological signal rather than technical artifacts. Together, these benchmarks evaluate both the breadth and precision of our model’s histopathology representations, covering frozen model’s performance, molecular inference, and multi-center robustness. The primary metric is the Robustness Index, which quantifies whether the nearest retrieved embedding corresponds to true biological signal or noise. This benchmark uses the concatenation of CLS token and average pooled patch tokens.

### 4.6 Compute

Training was performed on a cluster of 64 NVIDIA A100 GPUs across 8 nodes in a fully distributed, mixed-precision setting. Training required 5 days (i.e. 7680 GPU-hours), with over 80% of that time being devoted to Stage 1 training. GPU utilization was approximately 98% throughout training.

## 5 Declarations

### Model Availability

GenBio-PathFM is accessible at:

https://github.com/genbio-ai/genbio-pathfm

## A Supplemental Results

### A.1 Detailed Benchmark Results

The main paper presents figures with summary comparisons. In this section, we provide numerical results for each benchmark and its sub-tasks.

#### A.1.1 HEST

A detailed breakdown of the HEST benchmark results can be found in Table A1. In addition, Table A2 demonstrates that GenBio-PathFM can be jointly trained across all HEST tasks to improve performance.

**Table A1:**
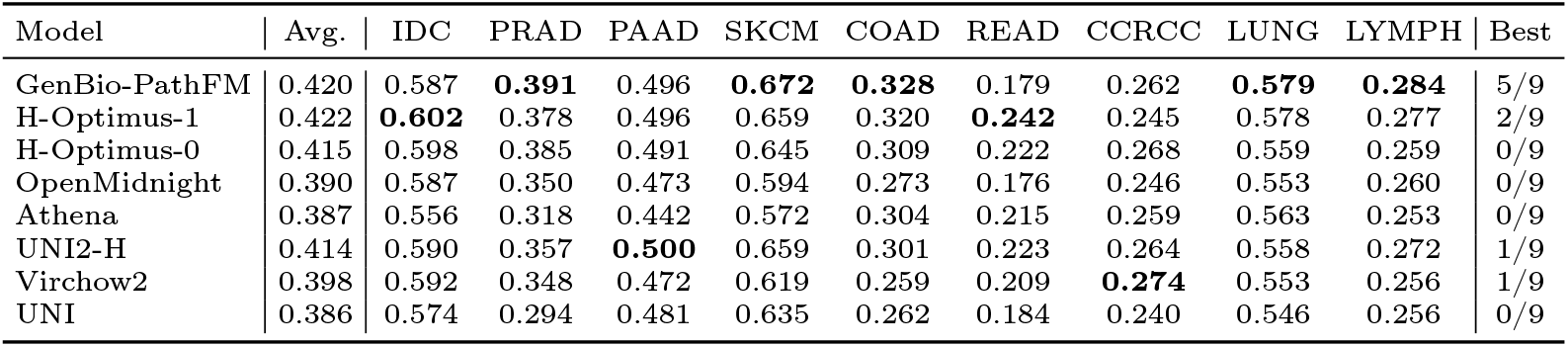
Detailed results on the HEST benchmark. For each sub-task, the highest score is bolded. The “Best” column counts the number of sub-tasks on which each model is the best performer.

| Model | Avg. | IDC | PRAD | PAAD | SKCM | COAD | READ | CCRCC | LUNG | LYMPH | Best |
| --- | --- | --- | --- | --- | --- | --- | --- | --- | --- | --- | --- |
| GenBio-PathFM | 0.420 | 0.587 | **0.391** | 0.496 | **0.672** | **0.328** | 0.179 | 0.262 | **0.579** | **0.284** | 5/9 |
| H-Optimus-1 | 0.422 | **0.602** | 0.378 | 0.496 | 0.659 | 0.320 | **0.242** | 0.245 | 0.578 | 0.277 | 2/9 |
| H-Optimus-0 | 0.415 | 0.598 | 0.385 | 0.491 | 0.645 | 0.309 | 0.222 | 0.268 | 0.559 | 0.259 | 0/9 |
| OpenMidnight | 0.390 | 0.587 | 0.350 | 0.473 | 0.594 | 0.273 | 0.176 | 0.246 | 0.553 | 0.260 | 0/9 |
| Athena | 0.387 | 0.556 | 0.318 | 0.442 | 0.572 | 0.304 | 0.215 | 0.259 | 0.563 | 0.253 | 0/9 |
| UNI2-H | 0.414 | 0.590 | 0.357 | **0.500** | 0.659 | 0.301 | 0.223 | 0.264 | 0.558 | 0.272 | 1/9 |
| Virchow2 | 0.398 | 0.592 | 0.348 | 0.472 | 0.619 | 0.259 | 0.209 | **0.274** | 0.553 | 0.256 | 1/9 |
| UNI | 0.386 | 0.574 | 0.294 | 0.481 | 0.635 | 0.262 | 0.184 | 0.240 | 0.546 | 0.256 | 0/9 |

**Table A2:** Comparison of GenBio-PathFM and a version of GenBio-PathFM that is trained using all of the HEST-bench training sets jointly. Performance increases on all sub-tasks.

| Model | Average | IDC | PRAD | PAAD | SKCM | COAD | READ | ccRCC | LUNG | LYMPH |
| --- | --- | --- | --- | --- | --- | --- | --- | --- | --- | --- |
| GenBio-PathFM | 0.420 | 0.587 | 0.391 | 0.496 | 0.672 | 0.328 | 0.179 | 0.262 | 0.579 | 0.284 |
| GenBio-PathFM (Joint) | 0.669 | 0.826 | 0.654 | 0.739 | 0.873 | 0.706 | 0.403 | 0.567 | 0.778 | 0.476 |
| % Change | +59% | +41% | +67% | +49% | +30% | +115% | +125% | +116% | +34% | +68% |

#### A.1.2 THUNDER

A detailed breakdown of the THUNDER benchmark results can be found in Table A3.

#### A.1.3 PathoROB

A detailed breakdown of the PathoROB benchmark results can be found in Table A4. Table A5 gives additional results on performance degradation due to distribution shift.

**Table A3:**
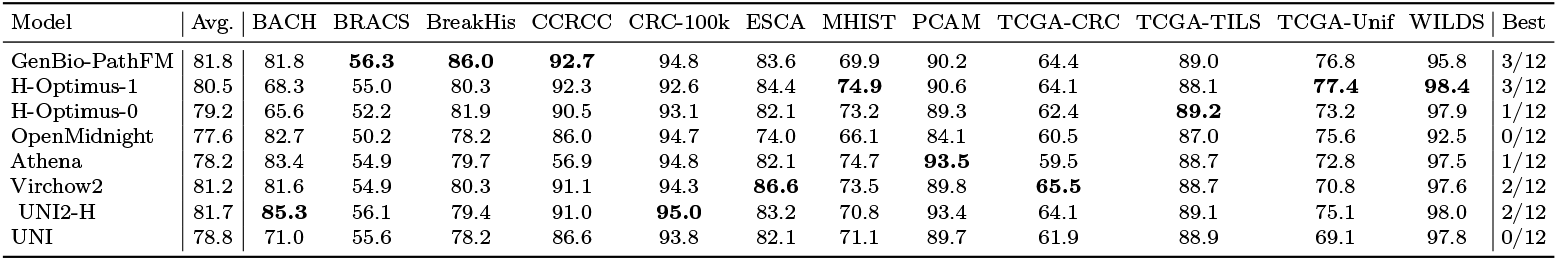
Detailed results on the THUNDER benchmark. For each sub-task, the highest score is bolded. The “Best” column counts the number of sub-tasks on which each model is the best performer.

**Table A4:** Results on the PathoROB benchmark, measured by robustness index (RI) and balanced accuracy (BACC). Higher values are better. The best performance in each column is bolded.

| Model | Camelyon |  | TCGA |  | Tolkach ESCA |  | Average |  |
| --- | --- | --- | --- | --- | --- | --- | --- | --- |
|  | RI | BACC | RI | BACC | RI | BACC | RI | BACC |
| GenBio-PathFM | **0.865** | 98.5 | 0.838 | 94.1 | **0.960** | **98.0** | **0.888** | 96.9 |
| H-Optimus-1 | 0.645 | **98.9** | **0.853** | **95.0** | 0.944 | 97.8 | 0.814 | **97.2** |
| H-Optimus-0 | 0.705 | 98.4 | 0.812 | 94.0 | 0.918 | 97.1 | 0.812 | 96.5 |
| OpenMidnight | 0.248 | 96.5 | 0.800 | 92.4 | 0.872 | 95.2 | 0.640 | 94.7 |
| Athena | 0.041 | 98.7 | 0.699 | 93.5 | 0.891 | 97.8 | 0.544 | 96.7 |
| Virchow2 | 0.806 | 98.8 | 0.822 | 92.8 | 0.955 | 97.7 | 0.861 | 96.4 |
| UNI2-H | 0.544 | 98.6 | 0.803 | 94.2 | 0.923 | 97.6 | 0.757 | 96.8 |
| UNI | 0.145 | 98.1 | 0.747 | 92.9 | 0.902 | 97.4 | 0.598 | 96.1 |

**Table A5:** Performance drop under in-distribution (ID) and out-of-distribution (OOD) shifts. More positive values are better. The best performance in each column is bolded.

|  | Camelyon |  | TCGA |  | Tolkach ESCA |  | Average |  |
| --- | --- | --- | --- | --- | --- | --- | --- | --- |
|  | ID | OOD | ID | OOD | ID | OOD | ID | OOD |
| GenBio-PathFM | **-2.3** | -0.6 | -0.9 | **0.8** | **0.2** | -0.1 | **-1.0** | **0.04** |
| H-Optimus-1 | -3.5 | -0.5 | -2.1 | -0.7 | -0.1 | -0.1 | -1.9 | -0.4 |
| H-Optimus-0 | -5.9 | **0.1** | -1.9 | -0.6 | -0.7 | -0.2 | -2.9 | -0.3 |
| OpenMidnight | -6.7 | -3.5 | -2.7 | -1.9 | -1.7 | -0.6 | -3.7 | -2.0 |
| Athena | -8.2 | -11.7 | -3.8 | 0.4 | -1.4 | -0.3 | -4.5 | -3.8 |
| Virchow2 | -2.9 | -0.4 | **-0.6** | -0.4 | -0.1 | **0.2** | -1.2 | -0.2 |
| UNI2-H | -4.6 | -1.3 | -2.6 | -2.6 | -0.4 | 0.04 | -2.6 | -1.3 |
| UNI | -9.6 | -1.4 | -2.1 | -2.5 | -0.1 | -0.02 | -4.3 | -1.3 |

### A.2 Performance-Robustness Tradeoff Results

Figure A1 shows the tradeoff between performance and robustness based on the PathoROB benchmark.

**Fig. A1:**
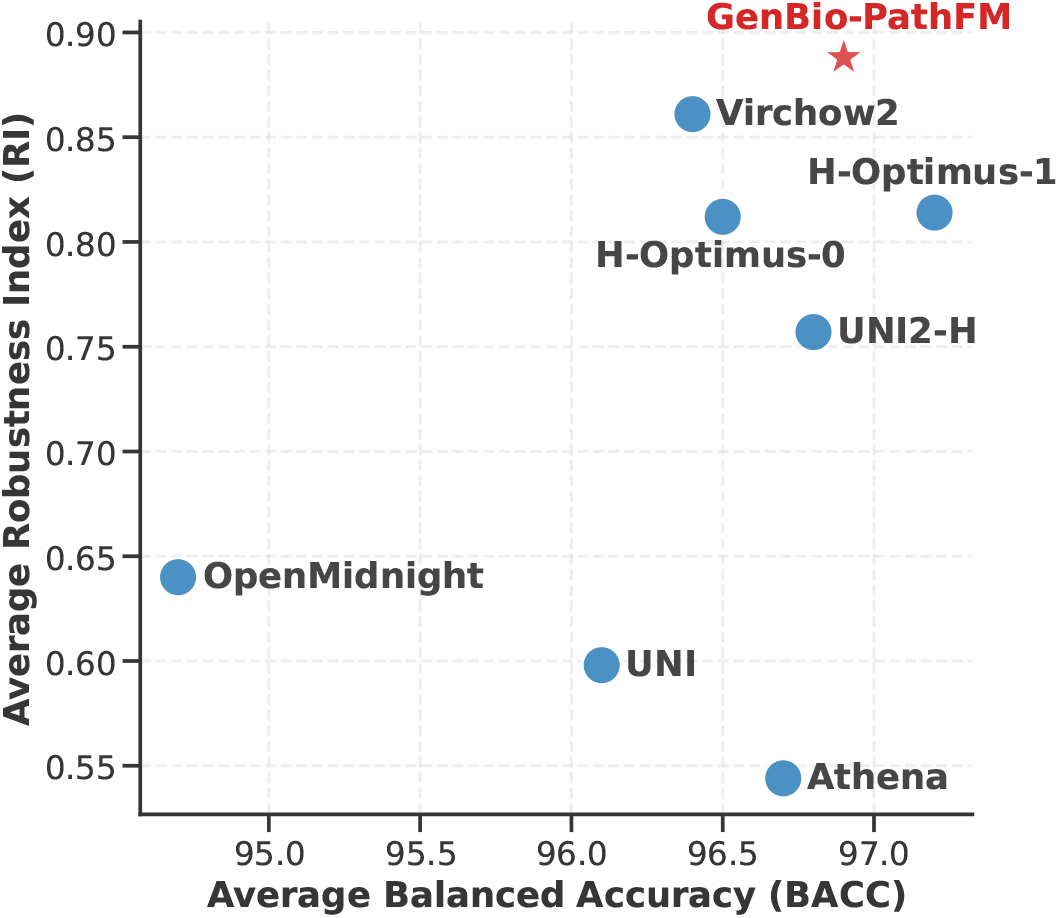
Performance-robustness tradeoff on the PathoROB benchmark.

### A.3 Ablation Studies

#### A.3.1 Pretraining Stage 1 vs. Stage 2

Table A6 compares the final version of GenBio-PathFM to the model after the first pretraining stage. We see that the second stage is an important contributor to performance.

**Table A6:** Ablation study results. Note that one task (MHIST) is responsible for the drop in performance on the THUNDER benchmark.

| Model Configuration | PathoROB | HEST | THUNDER | THUNDER (w/o MHIST) |
| --- | --- | --- | --- | --- |
| GenBio-PathFM (Stage 1 Only) | 0.838 | 0.410 | **82.1** | 82.2 |
| GenBio-PathFM | **0.888** | **0.420** | 81.8 | **82.9** |

#### A.3.2 Pretraining Stage 2 Design Choices

To further dissect the contributions of our novel JEDI Stage 2 approach, Table A7 details the impact of specific design choices on the HEST benchmark.

Starting from a baseline that applies a 75% random masking ratio and predicts only these masked regions, we iteratively introduced new components to the architecture. Reducing the masking ratio to 50%, adding the outpainting task (which requires the student to predict representations for regions beyond the visible crop), and adjusting the cropping parameters provided incremental benefits. We found that two key drivers were responsible for the substantial performance leaps.

The first critical driver was the addition of the visible region prediction objective. We speculate that this task serves to stabilize the training process. By explicitly forcing the student to match the teacher’s representations on unmasked areas, the model effectively retains the foundational representation learning capacity of the pretrained teacher. The student can then safely build upon this stable base through the more complex, spatially-aware tasks of masked region and outpainting prediction.

The second critical driver was explicitly omitting the loss on the CLS token. In standard Masked Autoencoder (MAE) pretraining, the objective is to reconstruct raw pixels for masked local patches, which naturally leaves the CLS token without a direct optimization target. In contrast, because our Stage 2 objective predicts representations in the embedding space of the teacher, we had the option to explicitly force the student’s CLS token to match the teacher’s CLS token. However, we found that removing this constraint yields superior results. This freedom may enable the CLS token to naturally aggregate the newly acquired fine-grained spatial information from the dense patch-level tasks, achieving a significant performance boost over the teacher’s original baseline. A closely related phenomenon was recently observed by [30], who evaluated the CLS token against average-pooled patch tokens in MAE-pretrained architectures. They found that even without an explicit loss applied to it, the CLS token still outperforms average pooling, suggesting that leaving the CLS token unconstrained in masked image modeling allows it to effectively form superior global representations.

### A.4 Qualitative Comparison of Tile Sampling Strategies

We show examples of tiles sampled randomly or through our curation approach in Figure A2.

**Table A7:** Ablation Study for JEDI Stage 2 Pretraining. The baseline uses a 75% random masking ratio (similar to MAE but in latent space of frozen teacher) without advanced prediction objectives. Subsequent steps introduce visible region prediction, reduced masking, outpainting, removal of the CLS token loss, and adjusted cropping parameters. The Δ column shows the performance difference of each configuration compared to the Stage 1 baseline performance (0.410).

| Experiment Step | Masking Ratio | Visible Region Pred. | Outpainting | No CLS Loss | Cropping Params | HEST Perf. | $\Delta$ vs. Stage 1 |
| --- | --- | --- | --- | --- | --- | --- | --- |
| 1. Baseline | 75% | – | – | – | Scale [0.428, 1.0]<br>AR [0.9, 1.1] | 0.3916 | -0.0184 |
| 2. Add Visible Pred. | 75% | ✓ | – | – | Scale [0.428, 1.0]<br>AR [0.9, 1.1] | 0.4072 | -0.0028 |
| 3. Reduce Masking | 50% | ✓ | – | – | Scale [0.428, 1.0]<br>AR [0.9, 1.1] | 0.4077 | -0.0023 |
| 4. Add Outpainting | 50% | ✓ | ✓ | – | Scale [0.428, 1.0]<br>AR [0.9, 1.1] | 0.4097 | -0.0003 |
| 5. Remove CLS Loss | 50% | ✓ | ✓ | ✓ | Scale [0.428, 1.0]<br>AR [0.9, 1.1] | 0.4196 | +0.0096 |
| 6. Adjust Cropping Params | 50% | ✓ | ✓ | ✓ | Scale [0.7, 1.0]<br>AR [0.75, 1.33] | 0.4198 | +0.0098 |

**Fig. A2:**
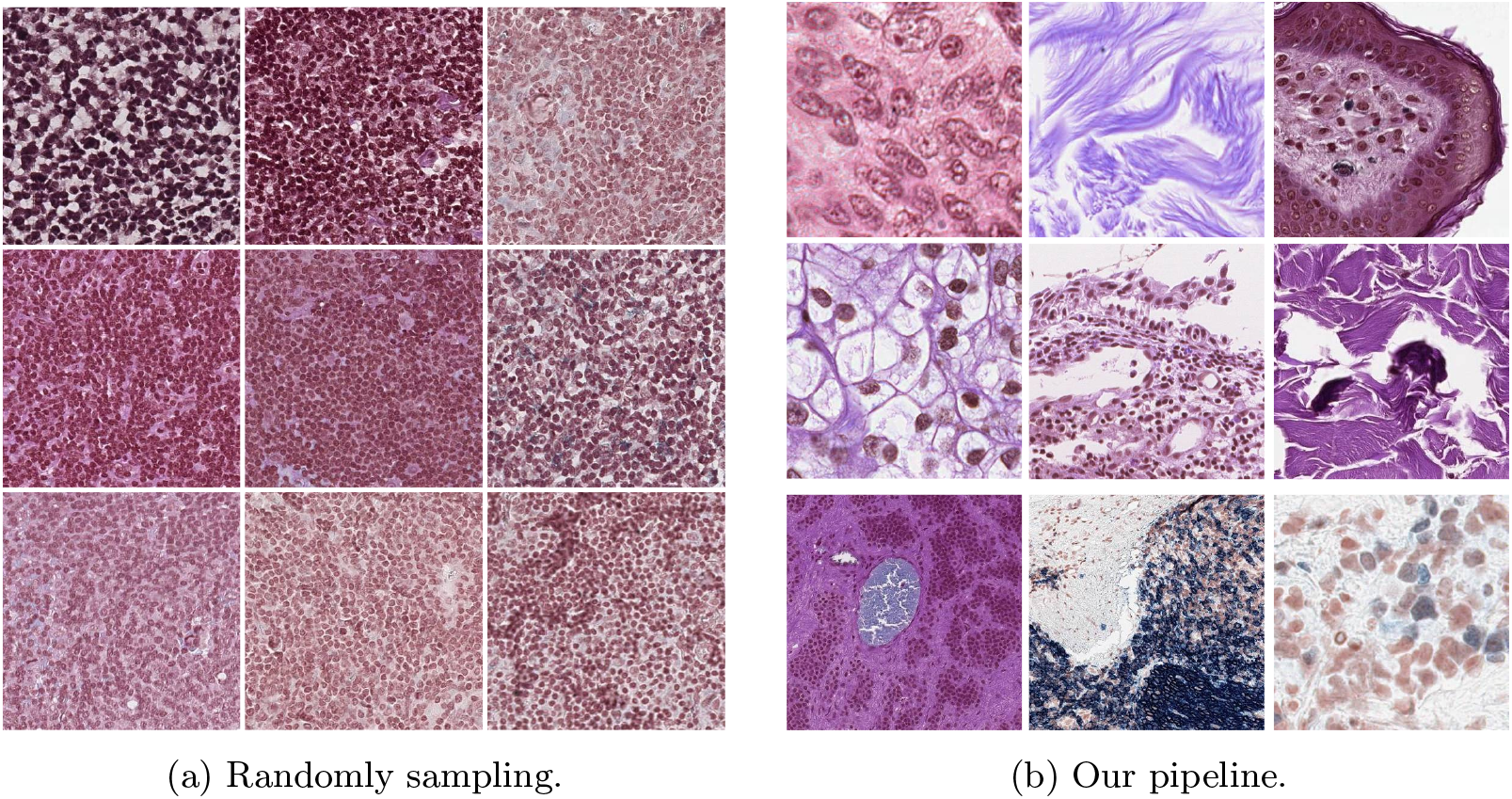
Comparison of tile sampling strategies. (a) Randomly sampled patches exhibit high redundancy with similar morphological patterns. (b) Tiles selected by our curation method demonstrate significantly greater morphological diversity.

## B Supplemental Methods

### B.1 Dataset Statistics

Table B8 describes the composition of our pretraining dataset. All of our WSIs are drawn from publicly available sources.

Table B9 compares the pretraining datasets for open-weight histopathology foundation models.

Tables B10, B11, B12, B13, and B14 give the sampling probabilities we used to build our pretraining datasets.

**Table B8:** Composition of our pretraining dataset in terms of the number of WSIs and the number of tiles extracted from those WSIs.

| Source | Approx. # WSIs | Approx. # Tiles |
| --- | --- | --- |
| HistAI [16] | 112k | 253M |
| TCGA [17] | 32k | 73M |
| GTEX [18] | 25k | 56M |
| REG [19] | 8k | 18M |
| Total | 177k | 400M |

**Table B9:** Comparison of pretraining datasets for open-weight histopathology foundation models.

| Model | # WSIs | Data Accessibility |
| --- | --- | --- |
| Virchow2 [3] | 3.1M | Private |
| H-Optimus-1 [13] | 1M | Private |
| H-Optimus-0 [31] | 500K | Private |
| UNI2-H [4] | 350K | Private |
| Athena [32] | 282K | Private |
| UNI [4] | 100K | Private |
| OpenMidnight [33] | 12K | Public |
| GenBio-PathFM | 177K | Public |

**Table B10:** Sampling probabilities for different magnification levels *m*.

| $m$ | Probability |
| --- | --- |
| 5× | 0.2 |
| 10× | 0.2 |
| 20× | 0.4 |
| 40× | 0.2 |
| **Total** | **1.00** |

**Table B11:** Conditional sampling probabilities for data sources given magnification level *m*.

| **Data Source** | $m \in \{5\times, 10\times, 20\times\}$ | $m = 40\times$ |
| --- | --- | --- |
| HistAI H&E | 0.40 | 0.06 |
| HistAI IHC | 0.12 | 0.03 |
| TCGA Diagnostic | 0.14 | 0.61 |
| TCGA Frozen | 0.12 | 0.30 |
| REG | 0.10 | 0.00 |
| GTE<sub>x</sub> | 0.12 | 0.00 |
| **Total** | **1.00** | **1.00** |

**Table B12:**
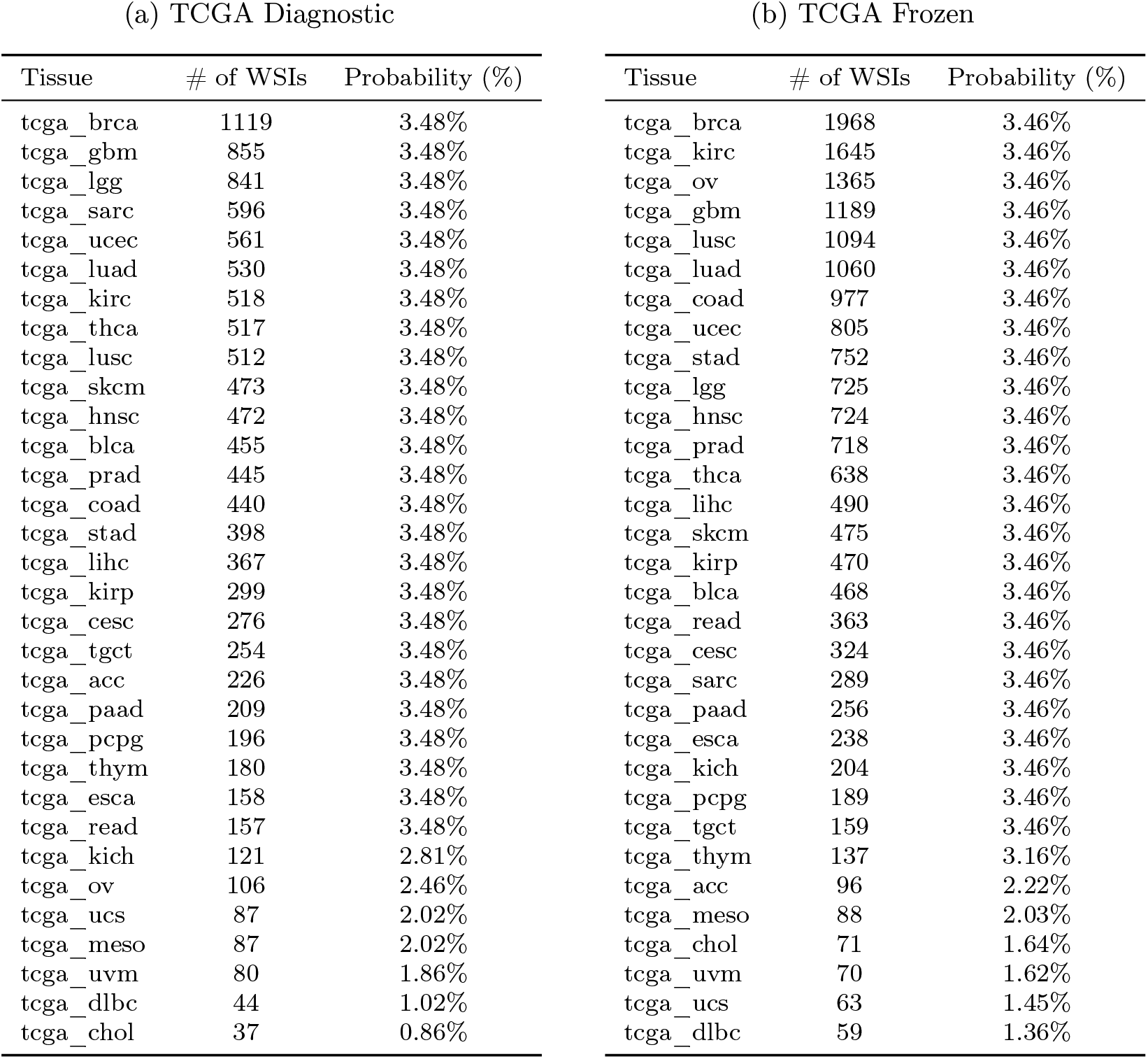
Conditional sampling probabilities for tissue types given data source and magnification level. These tables cover two of six data sources for magnifications *m* ∈ { 5×, 10×, 20× }. The other four data sources for these magnifications can be found in Table B13.

**Table B13:** Conditional sampling probabilities for tissue types given data source and magnification level. These tables cover four of six data sources for magnifications *m* ∈ { 5×, 10×, 20× }. The other two data sources for these magnifications can be found in Table B12.

| (a) HistAI H&E |  |  | (b) HistAI IHC |  |  |
| --- | --- | --- | --- | --- | --- |
| Tissue/Category | # of WSIs | Probability | Tissue/Category | # of WSIs | Probability |
| Skin | 49975 | 4.55% | Skin | 1386 | 4.55% |
| Breast | 5958 | 4.55% | Breast | 3275 | 4.55% |
| Thorax | 1145 | 4.55% | Thorax | 1191 | 4.55% |
| Hematologic | 1379 | 4.55% | Hematologic | 4392 | 4.55% |
| Endocrine | 594 | 4.55% | Endocrine | 275 | 4.55% |
| Head and neck | 718 | 4.55% | Head and neck | 468 | 4.55% |
| Gastrointestinal | 19952 | 22.73% | Gastrointestinal | 4548 | 22.73% |
| Gynecologic | 6747 | 22.73% | Gynecologic | 2040 | 22.73% |
| Genitourinary | 4730 | 22.73% | Genitourinary | 1111 | 22.73% |
| Others | 1231 | 4.55% | Others | 1578 | 4.55% |

**Table B13:** Conditional sampling probabilities for tissue types given data source and magnification level. These tables cover four of six data sources for magnifications $m \in \{5\times, 10\times, 20\times\}$ . The other two data sources for these magnifications can be found in Table B12.
| (c) REG |  |  | (d) GTex |  |  |
| --- | --- | --- | --- | --- | --- |
| Tissue | # of WSIs | Probability | Tissue | # of WSIs | Probability |
| Breast | 1919 | 22.59% | Esophagus | 2679 | 3.61% |
| Prostate | 1770 | 20.84% | Artery | 2467 | 3.61% |
| Stomach | 1477 | 17.39% | Skin | 1787 | 3.61% |
| Colon | 941 | 11.08% | Adipose | 1772 | 3.61% |
| Lung | 898 | 10.57% | Colon | 1720 | 3.61% |
| Bladder | 868 | 10.22% | Heart | 1360 | 3.61% |
| Cervix | 621 | 7.31% | Muscle | 989 | 3.61% |
|  |  |  | Nerve | 962 | 3.61% |
|  |  |  | Stomach | 929 | 3.61% |
|  |  |  | Thyroid | 893 | 3.61% |
|  |  |  | Breast | 886 | 3.61% |
|  |  |  | Spleen | 866 | 3.61% |
|  |  |  | Pancreas | 857 | 3.61% |
|  |  |  | Lung | 853 | 3.61% |
|  |  |  | Brain | 844 | 3.61% |
|  |  |  | Small Intestine | 791 | 3.61% |
|  |  |  | Adrenal Gland | 710 | 3.61% |
|  |  |  | Liver | 603 | 3.61% |
|  |  |  | Kidney | 598 | 3.61% |
|  |  |  | Prostate | 593 | 3.61% |
|  |  |  | Testis | 586 | 3.61% |
|  |  |  | Pituitary | 420 | 3.61% |
|  |  |  | Vagina | 272 | 3.61% |
|  |  |  | Ovary | 252 | 3.61% |
|  |  |  | Minor Salivary Gland | 245 | 3.61% |
|  |  |  | Uterus | 234 | 3.61% |
|  |  |  | Bladder | 129 | 3.11% |
|  |  |  | Cervix | 80 | 1.93% |
|  |  |  | Fallopian Tube | 43 | 1.04% |

**Table B14:**
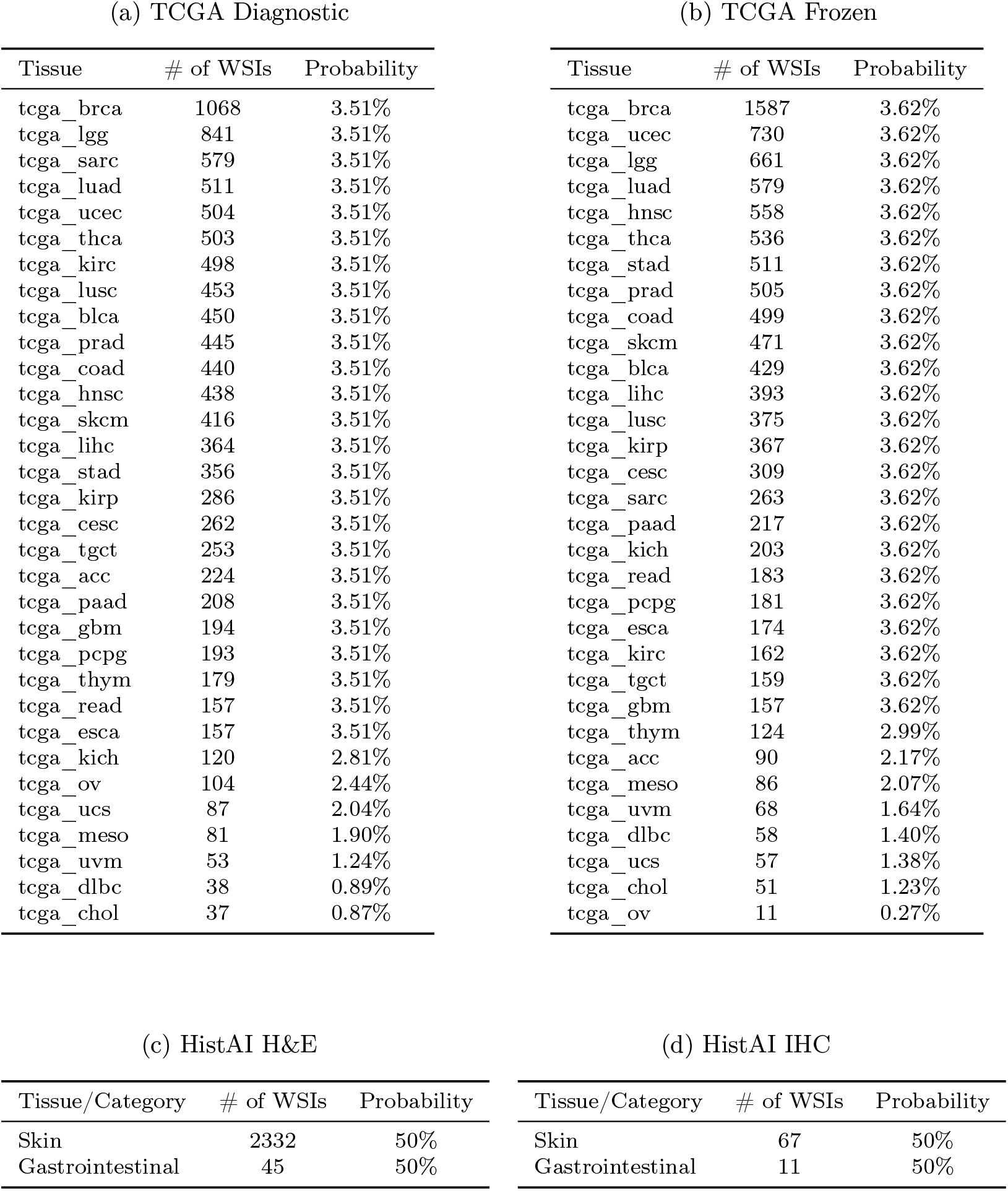
Conditional sampling probabilities for tissue types given data source and magnification level *m*, for *m* = 40×. See Table B12-B13 for analogous probabilities when *m* ∈ {5×, 10×, 20×}.

## References

[1] Marza, P., Fillioux, L., Boutaj, S., Mahatha, K., Desrosiers, C., Piantanida, P., Dolz, J., Christodoulidis, S., Vakalopoulou, M.: Thunder: Tile-level histopathology image understanding benchmark. In: The Thirty-ninth Annual Conference on Neural Information Processing Systems Datasets and Benchmarks Track

[2] Jaume, G., Doucet, P., Song, A.H., Lu, M.Y., Almagro-Pérez, C., Wagner, S.J., Vaidya, A.J., Chen, R.J., Williamson, D.F.K., Kim, A., Mahmood, F.: Hest-1k: A dataset for spatial transcriptomics and histology image analysis. In: Advances in Neural Information Processing Systems, vol. 37, pp. 53798–53833 (2024)

[3] Zimmermann, E., Vorontsov, E., Viret, J., Casson, A., Zelechowski, M., Shaikovski, G., Tenenholtz, N., Hall, J., Klimstra, D., Yousfi, R., et al.: Virchow2: Scaling self-supervised mixed magnification models in pathology. arXiv preprint 2408.00738 (2024)

[4] Chen, R.J., Ding, T., Lu, M.Y., Williamson, D.F., Jaume, G., Chen, B., Zhang, A., Shao, D., Song, A.H., Shaban, M., et al.: Towards a general-purpose foundation model for computational pathology. Nature Medicine (2024)

[5] Ma, J., Xu, Y., Zhou, F., Wang, Y., Jin, C., Guo, Z., Wu, J., Tang, O.K., Zhou, H., Wang, X., et al.: Pathbench: A comprehensive comparison benchmark for pathology foundation models towards precision oncology. arXiv preprint 2505.20202 (2025)

[6] Hosseini, M.S., Bejnordi, B.E., Trinh, V.Q.-H., Chan, L., Hasan, D., Li, X., Yang, S., Kim, T., Zhang, H., Wu, T., et al.: Computational pathology: a survey review and the way forward. Journal of Pathology Informatics 15, 100357 (2024)

[7] Caron, M., Touvron, H., Misra, I., Jégou, H., Mairal, J., Bojanowski, P., Joulin, A.: Emerging properties in self-supervised vision transformers. In: Proceedings of the IEEE/CVF International Conference on Computer Vision, pp. 9650–9660 (2021)

[8] Oquab, M., Darcet, T., Moutakanni, T., Vo, H.V., Szafraniec, M., Khalidov, V., Fernandez, P., Haziza, D., Massa, F., El-Nouby, A., et al.: Dinov2: Learning robust visual features without supervision. Transactions on Machine Learning Research

[9] Siméoni, O., Vo, H.V., Seitzer, M., Baldassarre, F., Oquab, M., Jose, C., Khalidov, V., Szafraniec, M., Yi, S., Ramamonjisoa, M., et al.: Dinov3. arXiv preprint 2508.10104 (2025)

[10] Kömen, J., Jong, E.D., Hense, J., Marienwald, H., Dippel, J., Naumann, P., Marcus, E., Ruff, L., Alber, M., Teuwen, J., et al.: Towards robust foundation models for digital pathology. arXiv preprint 2507.17845 (2025)

[11] Assran, M., Duval, Q., Misra, I., Bojanowski, P., Vincent, P., Rabbat, M., LeCun, Y., Ballas, N.: Self-supervised learning from images with a joint-embedding predictive architecture. In: Proceedings of the IEEE/CVF Conference on Computer Vision and Pattern Recognition, pp. 15619–15629 (2023)

[12] Li, X., Huang, C., Li, C.-L., Malach, E., Susskind, J., Thilak, V., Littwin, E.: Rethinking jepa: Compute-efficient video ssl with frozen teachers. International Conference on Learning Representations (2026)

[13] Bioptimus: H-optimus-1. https://huggingface.co/bioptimus/H-optimus-1

[14] Padigela, H., Nofallah, S., Chilaparasetti, A.N., Han, R., Walker, A., Shen, J., Shah, C., Martin, B., Sood, A., Miller, E., et al.: Pluto-4: Frontier pathology foundation models. arXiv preprint 2511.02826 (2025)

[15] Alber, M., Milbich, T., Carpen-Amarie, A., Tietz, S., Dippel, J., Muttenthaler, L., Cancer, B.P., Benetti, A., Korfiatis, P., Eulig, E., et al.: Atlas 2-foundation models for clinical deployment. arXiv preprint 2601.05148 (2026)

[16] Nechaev, D., Pchelnikov, A., Ivanova, E.: Histai: An open-source, largescale whole slide image dataset for computational pathology. arXiv preprint 2505.12120 (2025)

[17] Weinstein, J.N., Collisson, E.A., Mills, G.B., Shaw, K.R., Ozenberger, B.A., Ellrott, K., Shmulevich, I., Sander, C., Stuart, J.M.: The cancer genome atlas pan-cancer analysis project. Nature genetics 45(10), 1113–1120 (2013)

[18] Consortium, G., Ardlie, K.G., Deluca, D.S., Segrè, A.V., Sullivan, T.J., Young, T.R., Gelfand, E.T., Trowbridge, C.A., Maller, J.B., Tukiainen, T., et al.: The genotype-tissue expression (gtex) pilot analysis: multitissue gene regulation in humans. Science 348(6235), 648–660 (2015)

[19] REG2025 Grand Challenge. https://reg2025.grand-challenge.org/. Grand Challenge competition dataset (2025)

[20] Zhang, A., Jaume, G., Vaidya, A., Ding, T., Mahmood, F.: Accelerating data processing and benchmarking of ai models for pathology. arXiv preprint 2502.06750 (2025)

[21] Chen, B., Vincent-Cuaz, C., Schoenpflug, L.A., Madeira, M., Fournier, L., Subramanian, V., Andani, S., Ruiperez-Campillo, S., Vogt, J.E., Luisier, R., et al.: Revisiting automatic data curation for vision foundation models in digital pathology. In: International Conference on Medical Image Computing and Computer-Assisted Intervention, pp. 554–564 (2025). Springer

[22] Vo, H.V., Khalidov, V., Darcet, T., Moutakanni, T., Smetanin, N., Szafraniec, M., Touvron, H., Oquab, M., Joulin, A., Jegou, H., et al.: Automatic data curation for self-supervised learning: A clustering-based approach. Transactions on Machine Learning Research

[23] Radford, A., Kim, J.W., Hallacy, C., Ramesh, A., Goh, G., Agarwal, S., Sastry, G., Askell, A., Mishkin, P., Clark, J., et al.: Learning transferable visual models from natural language supervision. In: International Conference on Machine Learning, pp. 8748–8763 (2021). PmLR

[24] Dippel, J., Feulner, B., Winterhoff, T., Milbich, T., Tietz, S., Schallenberg, S., Dernbach, G., Kunft, A., Heinke, S., Eich, M.-L., et al.: Rudolfv: a foundation model by pathologists for pathologists. arXiv preprint 2401.04079 (2024)

[25] Su, J., Ahmed, M., Lu, Y., Pan, S., Bo, W., Liu, Y.: Roformer: Enhanced transformer with rotary position embedding. Neurocomputing 568, 127063 (2024)

[26] Sablayrolles, A., Douze, M., Schmid, C., Jégou, H.: Spreading vectors for similarity search. In: International Conference on Learning Representations (2019)

[27] Tellez, D., Litjens, G., Bándi, P., Bulten, W., Bokhorst, J.-M., Ciompi, F., Van Der Laak, J.: Quantifying the effects of data augmentation and stain color normalization in convolutional neural networks for computational pathology. Medical image analysis 58, 101544 (2019)

[28] Hu, S., Tu, Y., Han, X., He, C., Cui, G., Long, X., Zheng, Z., Fang, Y., Huang, Y., Zhao, W., et al.: Minicpm: Unveiling the potential of small language models with scalable training strategies. arXiv preprint 2404.06395 (2024)

[29] Loshchilov, I., Hutter, F.: Decoupled weight decay regularization. In: International Conference on Learning Representations (2019)

[30] Przewięźlikowski, M., Balestriero, R., Jasiński, W., Śmieja, M., Zieliński, B.: Beyond [cls]: Exploring the true potential of masked image modeling representations. In: Proceedings of the IEEE/CVF International Conference on Computer Vision, pp. 23442–23452 (2025)

[31] Saillard, C., Jenatton, R., Llinares-López, F., Mariet, Z., Cahané, D., Durand, E., Vert, J.-P.: H-optimus-0. https://github.com/bioptimus/releases/tree/main/models/h-optimus/v0

[32] Bosch, C., Wong, J.K., Paulikat, M., Zapukhlyak, M., Arora, B., Aichmüller-Ratnaparkhe, M., Baumann, J., Karn, S., Kamble, R., Karnik, S., et al.: Diversity over scale: Whole-slide image variety enables h&e foundation model training with fewer patches. Journal of Pathology Informatics, 100648 (2026)

[33] Kaplan, D., Grandhi, R.S., Lane, C., Warner, B., Abraham, T.M., Scotti, P.S.: How to Train a State-of-the-Art Pathology Foundation Model with $1.6k. https://sophont.med/blog/openmidnight

